# The NLRP3 inflammasome is essential for IL-18 production in a murine model of macrophage activation syndrome

**DOI:** 10.1101/2024.02.27.582284

**Authors:** Tara A. Gleeson, Christina Kaiser, Catherine B. Lawrence, David Brough, Stuart M. Allan, Jack P. Green

## Abstract

Hyperinflammatory disease is associated with an aberrant immune response resulting in cytokine storm. One such instance of hyperinflammatory disease is known as macrophage activation syndrome (MAS). The pathology of MAS can be characterised by significantly elevated serum levels of interleukin (IL)-18 and interferon (IFN)-γ. Given the role for IL-18 in MAS, we sought to establish the role of inflammasomes in the disease process. Using a murine model of CpG-DNA induced MAS, we discovered that the expression of the NLRP3 inflammasome was increased and correlated with IL-18 production. Inhibition of the NLRP3 inflammasome, or downstream caspase-1, prevented MAS-mediated upregulation of plasma IL-18 but interestingly did not alleviate key features of hyperinflammatory disease including hyperferritinaemia and splenomegaly. Furthermore IL-1 receptor blockade with IL-1Ra did not prevent the development of CpG-induced MAS, despite being clinically effective in the treatment of MAS. These data demonstrate that in the development of MAS, the NLRP3 inflammasome was essential for the elevation in plasma IL-18, a key cytokine in clinical cases of MAS, but was not a driving factor in the pathogenesis of CpG-induced MAS.

## Introduction

Cytokine storm syndromes (CSS) encompass a variety of disorders which present with hyperinflammation and multi-organ dysfunction characterised by excessive release of cytokines (hypercytokinemia) (*1*). CSS are generally defined by the underlying inflammation driving the cytokine storm response including infections (*2*), rheumatic diseases such as systemic juvenile idiopathic arthritis (SJIA) (*3*), adult-onset Still’s disease (AOSD) (*4*), systemic lupus erythematosus (SLE) (*5*), malignancy (*6*), immunotherapy (*7, 8*), or genetic defects such as primary haemophagocytic lymphohistiocytosis (pHLH) (*9, 10*). CSS are commonly termed as secondary hemophagocytic lymphohistiocytosis (sHLH) when they occur secondary to malignancy or infection, and more specifically in rheumatic disorders CSS is referred to as macrophage activation syndrome (MAS) (*11*). Crucially, if left untreated CSS can be lethal, highlighting the need to understand the biology behind CSS to provide effective therapies.

The characteristics of the cytokine storm depends on the causative factor, however in the case of MAS, several cytokines are proposed to contribute towards cytokine storm development, with the inflammasome derived cytokines interleukin (IL)-1β and IL-18 being implicated in disease pathogenesis (*12, 13*). IL-1β and IL-18 are produced as precursor proteins and are cleaved to generate biologically active forms by caspase-1 which is activated by inflammasomes (*14, 15*). Inflammasomes are multimolecular protein complexes containing a sensor pattern recognition receptor (PRR) such as NLRP3, NLRC4, AIM2 and NLRP1, the adaptor protein ASC (apoptosis associated speck-like protein containing a CARD), and the protease caspase-1 (*16-18*). Upon inflammasome formation caspase-1 is recruited leading to auto-proteolytic activation causing to two distinct events: 1) cleavage of pro-IL-1β and pro-IL-18 to their bioactive forms and 2) cleavage of the pore forming protein gasdermin-D (GSDMD) which allows for the release of mature IL-1β and IL-18 (*19*). The subsequent plasma membrane rupture is mediated by NINJ1 protein clustering which leads to the highly inflammatory form of cell death, known as pyroptosis (*20*). Both IL-1β and IL-18 signalling are intrinsically controlled by IL-1 receptor antagonist (IL-1Ra) and IL-18 binding protein (IL-18BP) respectively (*21, 22*).

Current understanding of MAS highlights two primary cytokines involved in the pathogenesis of the cytokine storm: interferon (IFN)γ and IL-18. Furthermore, IL-18 is understood to be a key driver of IFNγ production indicating that these cytokines are partaking in a feedback loop (*23, 24*). Clinically, IL-18 is used as an important marker of MAS (*25-27*). Circulating IL-18 levels are elevated in both patients with SJIA and AOSD (*28-30*) and are significantly increased during episodes of MAS (*13, 31*) and this significant elevation in IL-18 diagnostically distinguishes MAS flares from underlying rheumatic disease (*25*). Mouse models of MAS/HLH also present with increased levels of IL-18 (*25, 32*) and development of MAS is worsened in IL-18BP knockout mice (*32*). Treatment with recombinant IL-18BP (tadekinig alfa) has been shown to reduce symptoms of MAS in AOSD patients (*33*). Further, there are also proposed roles for IL-1 cytokines in MAS pathogenesis since IL-1Ra is currently used off label for the treatment of CSS, with patients responding well to high doses (*34-36*). Mouse models of hyperinflammatory disease also demonstrate an IL-18 signature in the blood, in particular a mouse model of MAS that is induced by repeated administration of cytosine guanine 1826 oligonucleotide (CpG)-DNA induces a phenotype similar to that observed in patients with MAS (*37*). Other models of hyperinflammatory disease rely more on IFNγ signalling rather than IL-18 signalling, more closely resembling pHLH (*38, 39*). The CpG-induced MAS model has been used to uncover the dynamics of IL-18/IFNγ signalling in disease (*32*) (*40*) (*41*). Despite evidence for the involvement of IL-1 and IL-18 cytokines in MAS pathogenesis, the mechanisms driving MAS remain unclear. Presently, the NLRC4 inflammasome has been implicated in one instance of MAS, known as NLRC4-MAS, where gain-of-function mutations in NLRC4 drive MAS pathogenesis (*42-44*). However, the mechanisms promoting other instances of CSS and MAS remain unclear and the role of inflammasomes in these has yet to be fully elucidated.

In this study, we investigated the role of inflammasomes and IL-1 cytokines in hyperinflammation using a mouse model of CpG DNA-induced MAS. Here, we show that the NLRP3 inflammasome is upregulated in the development of MAS, with tissues displaying elevated levels of NLRP3, caspase-1 and IL-18 following induction of hyperinflammation. However, pharmacological inhibition of either the NLRP3 inflammasome, caspase-1 or the IL-1 receptor did not prevent development of MAS symptoms, despite a reduction in plasma IL-18 levels following NLRP3 or caspase-1 inhibition. Our data suggest that whilst the NLRP3 inflammasome is responsible for increased circulating IL-18 in MAS, it is not responsible for the development of other features of MAS pathogenesis in CpG DNA-induced MAS, such as hyperferritinaemia and splenomegaly. These data suggest that alternative mechanisms are responsible for the splenomegaly, hypercytokinemia and organ dysfunction in MAS.

## Methods

### Animals

Male 8–12-week-old C57BL/6J mice (Charles River Laboratories, UK) were used in all experiments. Animals were housed in individually ventilated cages with temperature and humidity maintained between 20–24°C and 45 to 65%, respectively. Animals were housed in a room with a 12 h light-dark cycle. All animal experiments were carried out under the authority of a UK Home Office Project Licence and reported according to the ARRIVE guidelines for experiments involving animals (*45*).

### Induction of CpG DNA-induced MAS

Mice were treated with CpG ODN 1826 oligodeoxynucleotide (synthesised by Integrated DNA technologies (IDT); 5’-T^*^C^*^C^*^A^*^T^*^G^*^A^*^C^*^G^*^T^*^T^*^C^*^C^*^T^*^G^*^A^*^C^*^G^*^T^*^T-3’, where ^*^ indicates a phosphorothioate modification) (2 mg/kg) or vehicle control (sterile phosphate buffered saline; PBS, 10 µL/g) five times over the course of ten days, as described previously (*37*). Mice received CpG or PBS by intraperitoneal (I.P) injection on day 0, 2, 4, 7 and 9 of the protocol. Animals were weighed daily, and then culled on day 10 (at 24 hour-post injection), or at the indicated time-point for time course experiments. The following treatments were given (I.P. unless stated): MCC950 (50 mg/kg in PBS, (*46*)) (CP-456773 sodium salt, Sigma Aldrich) at the same time as CpG; vehicle controls received 10 µL/g PBS; VX765 (Belnacasan) (100mg/kg in 5% (v/v) DMSO in PBS, (*47*)) (273404-37-8, Tocris Bioscience) injected daily; vehicle controls received 10 µL/g 5% (v/v) DMSO in PBS; recombinant IL-1Ra (100mg/kg in placebo (638 mM polysorbate 80, 5.2 mM sodium citrate, 112 mM sodium chloride, 45.4 µM disodium EDTA, dH2O, pH 6.5)) (Anakinra, Sobi) administered twice daily (*48*); high dose IL-1Ra recommended for clinical treatment of CSS (*49, 50*) by sub-cutaneous (S.C) injection; vehicle controls received 10 µL/g placebo. Researchers were blinded to treatment for the duration of the experiment.

### Tissue Collection

On day 10 mice were deeply anaesthetised with 2.5% isoflurane (Isofane, Henry Schein) in 33% O_2_ and 67% NO_2_ and blood taken via cardiac puncture. Blood was spun at 1,500xg for 15 min, plasma removed and spun again at 14,000xg for 3 min. Plasma was aliquoted and stored at -80°C for analysis. Following cardiac puncture, mice were perfused with PBS, spleen was removed, and weight recorded. Spleen was then divided for further analysis. Liver was removed and dissected for further analysis. A proportion of liver and spleen were snap frozen for western blot analysis, and another portion was drop fixed in 4% paraformaldehyde (PFA) and then embedded in paraffin.

### ELISA

Sandwich ELISA was used to establish plasma concentrations of IL-18 (Invitrogen, Thermofisher, BMS618-3) and ferritin (Abcam, ab157713). ELISA protocol was carried out according to manufacturer’s instructions. ELISA was read at 450-570nm according to manufacturer’s instructions.

### Multiplex cytokine analysis assay

The concentration of IFNγ, TNF, IL-10, and IL-6 were measured by LEGENDplex flow-based 13-plex mouse inflammation panel kit from Biolegend (740446) according to manufacturer’s instructions. LEGENDplex results obtained using BD FACSVerse (BD Biosciences).

### Tissue Homogenisation

Livers and spleens were snap frozen on dry ice immediately following isolation and stored at -80°C prior to homogenisation. Spleens were homogenised in RIPA lysis buffer (150mM sodium chloride, 1.0% (v/v) NP-40, 0.5% (w/v) sodium deoxycholate, 0.1% (w/v) sodium dodecyl sulfate, 50mM Tris pH 8.0, dH2O) containing protease inhibitor cocktail (Sigma, 539131-10). Livers were homogenised in NP-40 lysis buffer (0.5% (v/v) NP-40, 150mM sodium chloride, 2mM EDTA, 50mM Tris pH 8.0, dH2O) containing protease inhibitor cocktail (Sigma, 539131-10). Protein concentration of homogenates was determined using BCA (bicinchoninic) protein assay kit (ThermoFisher, 23225). BCA assay was used to determine protein concentration of each sample and samples were used at a concentration of 175µg for western blotting.

### Western Blotting

Laemmli buffer (5x) was added to samples and boiled at 95°C for 10 min before resolving by SDS-PAGE. Resolved gels were transferred onto nitrocellulose or PVDF membranes using a Trans-Blot® Turbo Transfer™ System (Bio-Rad). Membranes were blocked in 5% (w/v) milk in PBS 0.1% Tween-20 (PBST) for 1 h at room temperature. Membranes were washed with PBST and incubated overnight with rabbit-anti-mouse IL-18 (1/1000 dilution; E9P50, Cell Signalling Technology, 57058), rabbit-anti-mouse capase-1 p10 (1/1000 dilution; EPR16883, Abcam, ab179515), mouse-anti-mouse NLRP3 (1/1000 dilution; Cryo2, Adipogen, AG-20B-0014), rabbit-anti-mouse gasdermin-D (1/1000 dilution; EPR19828, Abcam, ab209845), or goat-anti-mouse IL-1β (1/800 dilution; R&D Systems, AF-401-NA), primary antibodies were diluted in 5% (w/v) BSA in PBST. The membranes were washed and incubated at room temperature for 1 h with rabbit anti-mouse IgG (Agilent, P026002-2) or goat anti-rabbit IgG (Agilent, P044801-2), secondary antibodies diluted 1/1000 in 5% (w/v) BSA in PBST. Proteins were then visualized with Amersham ECL Prime Western Blotting Detection Reagent (Cytiva, RPN2236) and G:BOX (Syngene) and Genesys software. β-Actin (1/20000 dilution 5% (w/v) BSA in PBST; Sigma, A3854) was used as a loading control.

### Histology

Spleen sections (5 µm, cut using paraffin rotary microtome (Leica)) were stained with Haematoxylin (ThermoFisher) and Eosin Y (ThermoFisher) (H&E). Coverslips were applied using DPX mountant (Sigma, 06522). Spleen sections were stained for iron with Perls Prussian Blue Stain Kit (Abcam, 65692) according to manufacturer’s instructions. Sections were scanned on SlideScanner (3D Histech Panoramic P250) and analysed on CaseViewer (3Dhistech Ltd.).

### Statistical Analysis

Data were analysed using GraphPad PRISM 9 software (GraphPad Software Inc. CA, USA). Results are presented as mean + SEM. Equal variance and normality were assessed using the Shapiro-Wilk test. To compare two data sets (PBS vs CpG), an unpaired Student’s *t*-test was chosen. For data with two factors (PBS/CpG ± vehicle/treatment), a one-way ANOVA with Tukey’s multiple comparison test was performed. Non-parametric data was transformed before statistical analysis. Accepted levels of significance ^*^p<0.05, ^**^p<0.01, ^***^p<0.001, ^****^p<0.0001. Studies conducted on groups of 3 or 5 animals. N represents an individual animal.

## Results

### Repeated TLR9 stimulation results in the initiation of hyperinflammatory disease and upregulates the inflammasome

We used a previously described murine model of MAS, where repeated intraperitoneal administration of the TLR9 agonist CpG-DNA (ODN 1826) induces features of hyperinflammatory disease, similar to those observed in patients with MAS (*37*). We administered CpG-DNA (ODN 1826) at 2 mg/kg five times over the course of the 10 days (Figure 1A). Matching previous reports, CpG-treated mice transiently lost weight following the administration of CpG-DNA (Figure 1B). CpG treatment induced significant splenomegaly (Figure 1C, D). Further, CpG-DNA administration resulted in a significant increase in plasma ferritin levels (Figure 1E), a marker of inflammatory disease (*51*), and a significant increase in plasma cytokines, emulating cytokine storm associated with hyperinflammatory disease. CpG-DNA treated mice had significantly elevated levels of plasma IFNγ, IL-18, IL-6, IL-10 and TNF compared to PBS controls (Figure 1F-J), but plasma levels of IL-1β and IL-1α were below the limit of detection (data not shown). The CpG-DNA induced mouse model of MAS has been well established as an appropriate model for “subclinical” MAS as it recapitulates a number of the pathologies associated with the disease and is worsened by removal of the endogenous regulator of IL-18 signalling, IL-18BP (*32*). However, the source of IL-18 in this model has not been previously reported. Therefore, we assessed if expression of components of the inflammasome were increased following CpG-induced hyperinflammation. Homogenised spleens from mice with repeated CpG injections had enhanced expression of inflammasome components including NLRP3, caspase-1, gasdermin-D, pro-IL-1β and pro-IL-18 (Figure 1K). These data show that repeated administration of CpG-DNA produces markers of hyperinflammation and suggests that inflammasome signalling is upregulated in the development of MAS. This provides the first evidence that CpG-induced MAS leads to upregulation of inflammasome components, indicating a possible role for inflammasomes in disease pathogenesis.

**Figure 1:**
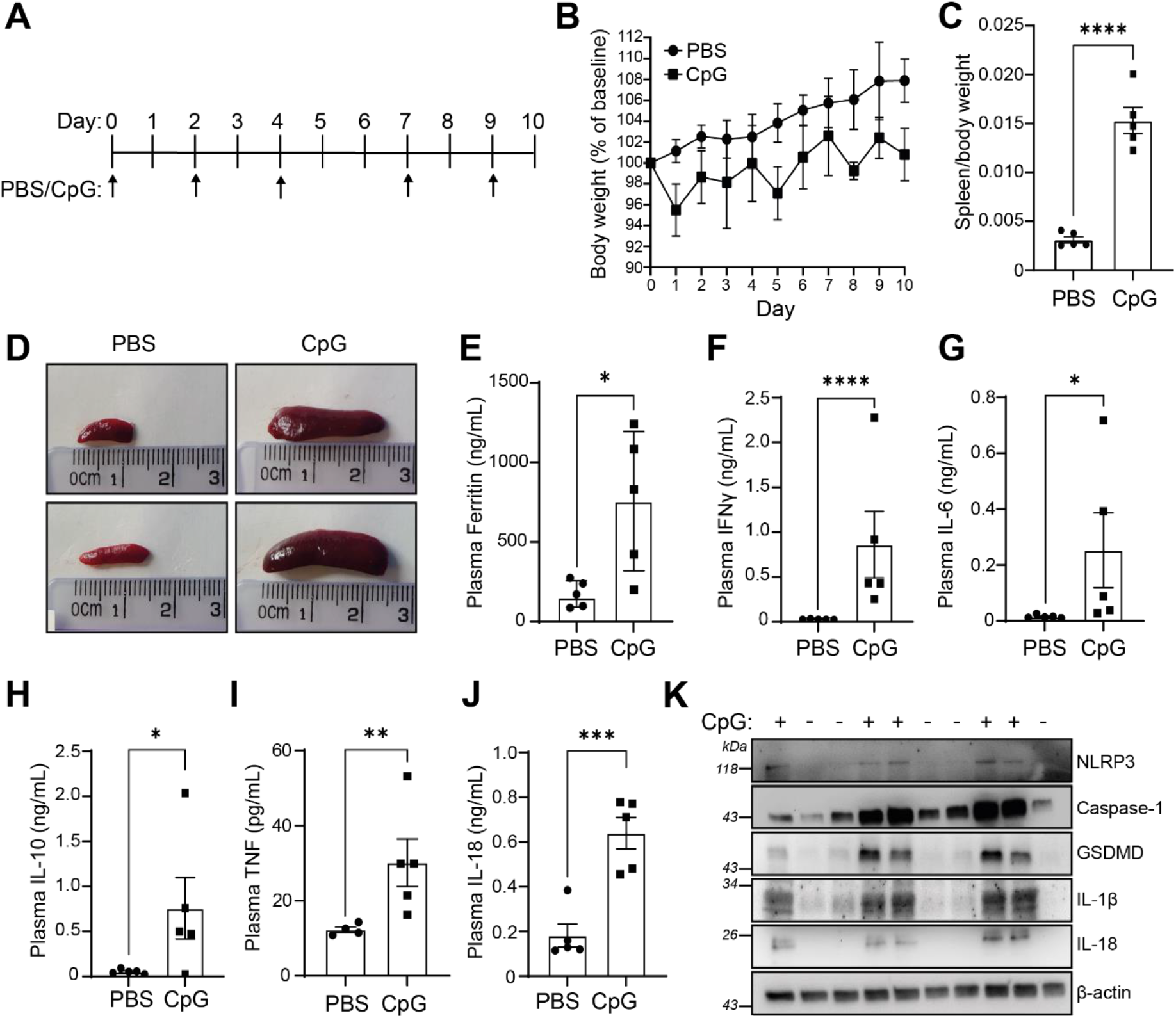
Repeated administration of CpG-DNA induced characteristics of hyperinflammatory disease and inflammasome signalling. **(A)** Mice were treated over a 10-day period with CpG-DNA (ODN 1826, 2mg/kg), or vehicle (PBS), by intraperitoneal injection on days 0, 2, 4, 7 and 9. **(B)** Animal weight was measured daily over the course of the study. Animal weights presented as a percentage of weight on day 0. Animals were sacrificed on day 10 and hyperinflammatory readouts were recorded. **(C)** Splenic weight normalised to body weight from CpG-DNA (CpG) or PBS injected mice (n=5). **(D)** Representative images from (C). **(E)** Plasma levels of ferritin (n=5). **(F-J)** Plasma concentration of IFNγ (F), IL-6 (G), IL-10 (H) and TNF (I), IL-18 (J) (n=5). **(K)** Western blot of homogenised spleens from PBS or CpG treated animals were blotted for inflammasome components NLRP3, caspase-1, GSDMD and IL-1β (n=5). Data represent the mean ± SEM. ^*^, P<0.05, ^**^, P<0.01, ^***^, P<0.001, ^****^, P<0.0001 determined by an unpaired Student’s *t*-test.

We also wanted to assess the temporal dynamics of inflammasome upregulation to further dissect the pathogenesis of disease and ensure that day 10 was the optimal day for analysis of inflammasome involvement in MAS. Therefore, we assessed the response to a single dose of CpG-DNA at 6 and 24 h, as well as the effects after 2, 3, 4 or 5 repeated doses (Supplemental Figure 1A). Splenomegaly was proportional to number of doses received (Supplemental Figure 1B). CpG-DNA caused an initial hyperferritinaemic response within 24 h of administration, which remained elevated after dose 2, 3 and 4 until increasing further following the 5^th^ dose (Supplemental Figure 1C). We observed different induction kinetics between cytokines. Plasma IFNγ, IL-6 and TNF peaked 24 h after CpG administration before decreasing over time, although the concentration of IFNγ remained elevated at all time points compared to naïve animals (Supplemental Figure 1D, E and G). Conversely, plasma IL-18 levels exhibited an initial acute elevation, which resolved before a large and sustained increase at later stages of MAS development following 3 injections (Supplemental Figure 1H), the same was seen with IL-10 (Supplemental Figure 1F). These data indicate that the repeat CpG-DNA model presents with an initial acute inflammatory response, followed by the development of a more consistent hyperinflammatory phenotype that recapitulates what is observed clinically with hyperinflammatory diseases, with inflammasome activation occurring later in the disease time course. Both liver and spleen homogenates indicated a dose dependent increase in NLRP3, pro-caspase-1 and pro-IL-18 expression (Supplemental Figure 1I, J). Histological analysis of the liver revealed that CpG-treated mice had a marked reduction in iron (Fe^3+^) in the liver over time, correlating with CpG-DNA doses (Supplemental Figure 1K). Perl’s Prussian blue staining correlated with ferritin concentration in the plasma, indicative of iron sequestering and inflammation. These data suggest that the upregulation of the NLRP3 inflammasome occurred later in the pathogenesis of CpG-induced MAS and that inflammasome expression coincided with plasma IL-18 activity.

### The NLRP3 inflammasome is dispensable for the development of CpG-induced MAS

To better understand the role of inflammasomes in CpG-induced MAS, and since we had observed an upregulation of the NLRP3 inflammasome following repeated CpG injections, we then tested if the NLRP3 inflammasome was critical for the development of CpG-induced hyperinflammation. To do this, mice were treated with the NLRP3 specific inhibitor MCC950 (*46*) (50 mg/kg, I.P.) in tandem with CpG-DNA or PBS every 2 days over a 10-day period (Figure 2A). As before, CpG-DNA treatment induced significant splenomegaly, but this was not reduced following MCC950 co-treatment (Figure 2B, C). Further, CpG-induced hyperferritinaemia was unaffected by MCC950 treatment (Figure 2D). Examination of splenic architecture revealed a disruption in normal red and white pulp morphology in CpG treated animals which persisted in MCC950-treated animals (Figure 2E). We then assessed if inflammasomes were contributing to the development of the cytokine storm in CpG-induced hyperinflammation by analysing plasma cytokines in mice treated with repeated CpG and NLRP3 inhibitors (Figure 2F-J). MCC950 treatment significantly reduced CpG-induced plasma IL-18 to similar levels as seen in PBS-injected animals (Figure 2J), suggesting that the NLRP3 inflammasome is responsible for elevated plasma IL-18. However, MCC950 treatment did not significantly alter production of IFNγ, IL-6, IL-10, or TNF in response to repeated CpG (Figure 2F-I), though there was a trend toward an increase in plasma IFNγ concentrations (P=0.2810) (Figure 2F). These results demonstrate that inhibition of the NLRP3 inflammasome is not sufficient to prevent onset of CpG-induced hyperinflammatory disease.

**Figure 2:**
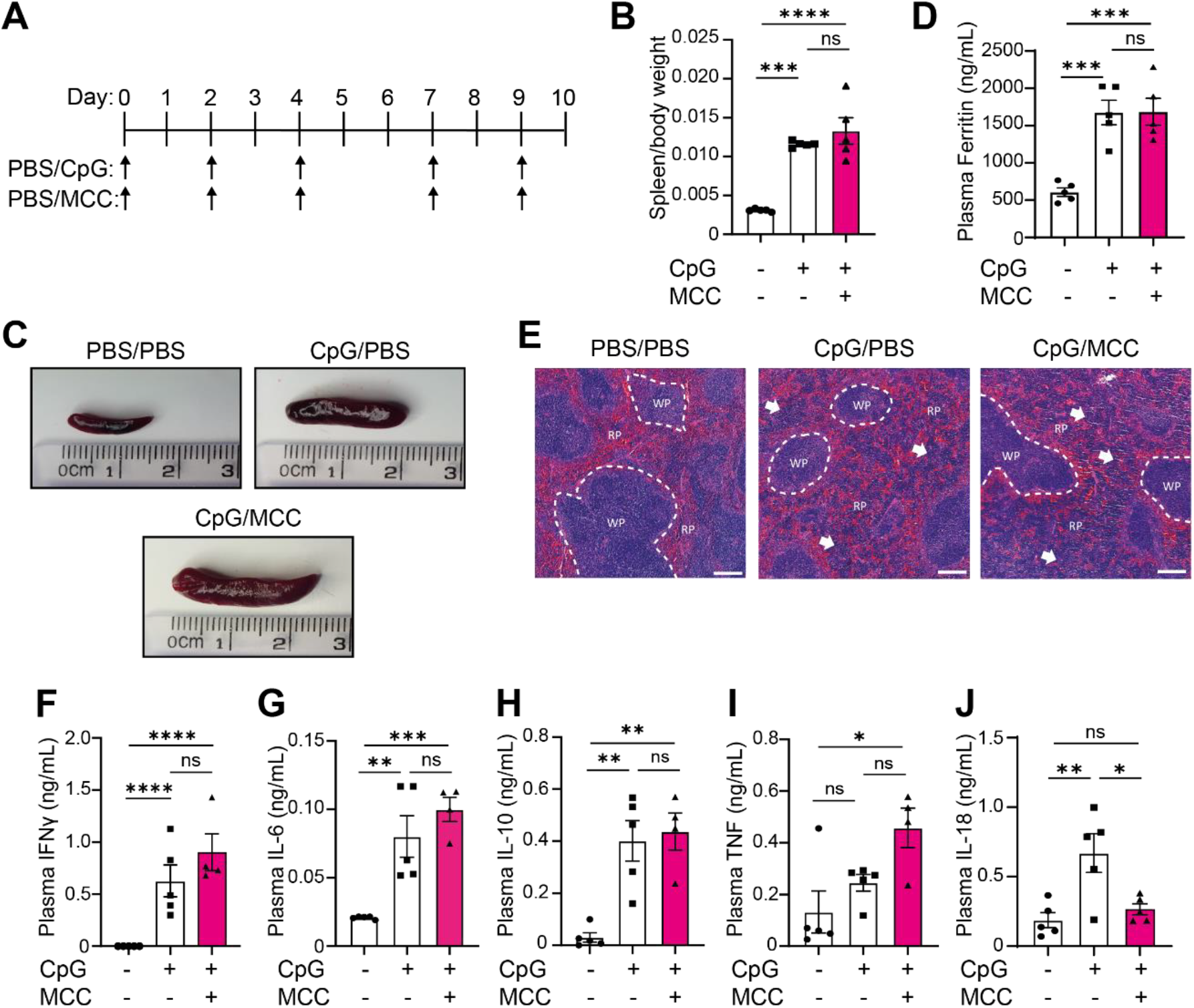
The NLRP3 inflammasome is dispensable for the development of CpG-induced hyperinflammation. **(A)** Mice received 5 doses of CpG-DNA (CpG, ODN 1826, 2 mg/kg), along with vehicle (PBS) or MCC950 (MCC, 50 mg/kg). **(B)** Splenic weight normalised to body weight from CpG-DNA (CpG) or PBS injected mice (n=5). **(C)** Representative images from **(B). (D)** Plasma levels of ferritin (n=5). **(E)** H&E staining of spleen in mice treated with PBS/PBS, CpG/PBS or CpG/MCC. RP = red pulp, WP = white pulp. White arrows denote changes to normal splenic architecture and perturbations to red pulp. Scale bar represents 200 µm. **(F-J)** Plasma concentration of IFNγ (F), IL-6 (G), IL-10 (H) and TNF (I), IL-18 (J) (n=5). Data represent the mean ± SEM. ^*^, P<0.05, ^**^, P<0.01, ^***^, P<0.001, ^****^, P<0.0001 determined by one-way ANOVA with Tukey’s multiple comparisons test.

### Inflammasome activation is not required to drive CpG-induced MAS

Since inhibition of the NLRP3 inflammasome did not prevent hyperinflammatory disease, we then questioned if alternative inflammasomes could contribute towards the hyperinflammatory state. To test this, we used a caspase-1 inhibitor, VX765 (Belnacasan). As mentioned previously, inflammasome activation leads to recruitment and cleavage of caspase-1 (*16*) (*17*) (*18*), meaning that inhibition of caspase-1 activity results in pan-inflammasome inhibition (such as NLRP3, AIM2, NLRC4, NLRP1). To decipher the impact of inflammasome activation on MAS pathogenesis, we repeated our 10-day model of CpG-induced hyperinflammation with mice treated daily with the caspase-1 inhibitor VX-765 (*47*) (100 mg/kg, I.P.) (Figure 3A). Caspase-1 inhibition with VX765 did not affect development of splenomegaly (Figure 3B, C), hyperferritinaemia (Figure 3D), or prevent perturbations in splenic architecture (Figure 3E), similar to what was observed with MCC950 treatment. Likewise, there were no significant changes to plasma levels of IFNγ, IL-6, IL-10 or TNF (Figure 3F-I), but a significant reduction in IL-18 was observed (Figure 3J) (P=0.0007), along with a non-significant trend to increased plasma IFNγ concentrations (P=0.0596) (Figure 3F). These data indicate that inflammasomes are not essential in CpG-induced splenomegaly, elevated plasma ferritin, splenic tissue disruption or cytokine storm, apart from plasma IL-18 concentrations. Further, since caspase-1 inhibition exhibited the same phenotype as NLRP3 inhibition, this suggests that only the NLRP3 inflammasome is responsible for enhanced plasma IL-18 in CpG-induced MAS.

**Figure 3:**
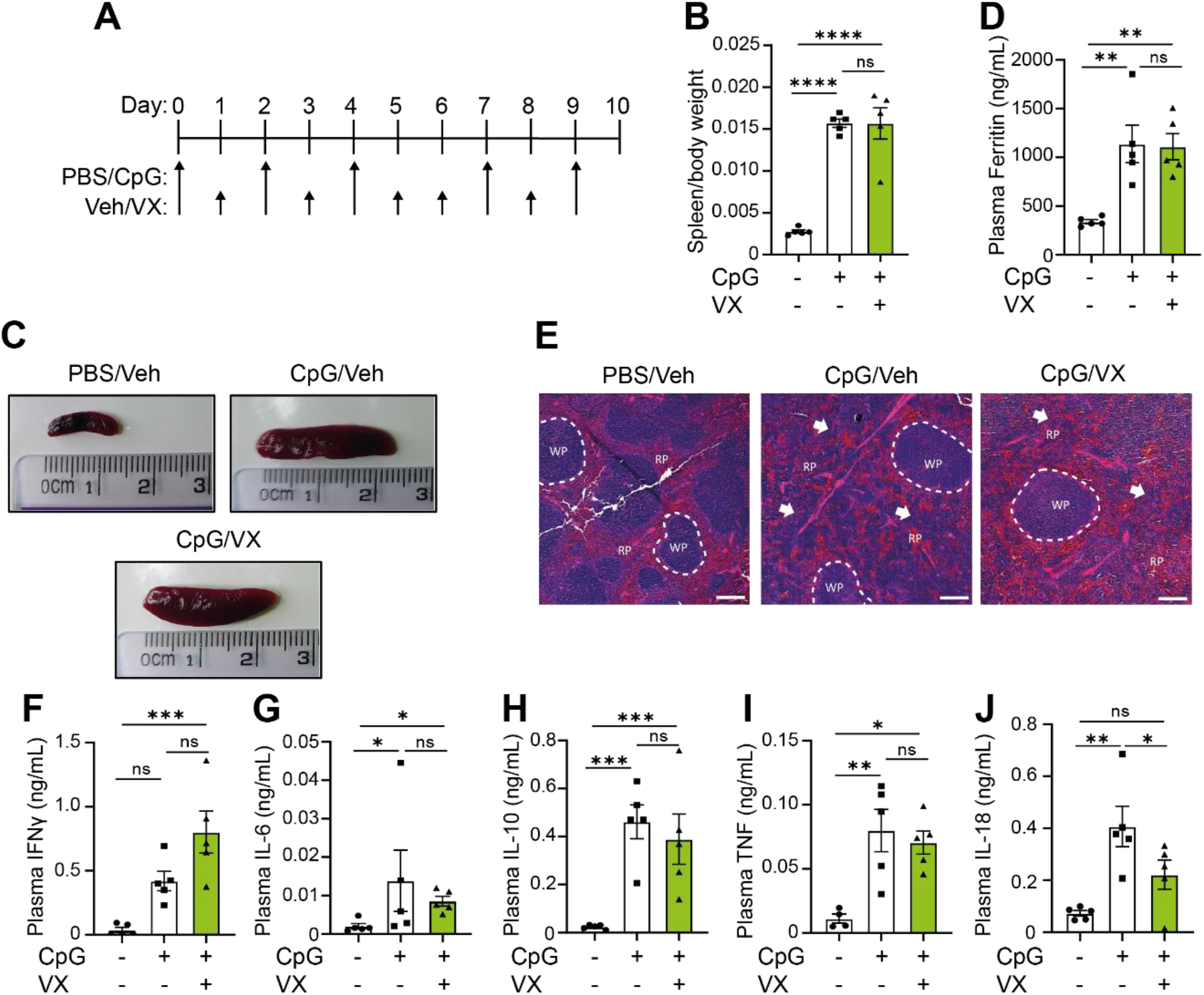
Canonical inflammasome activation is dispensable for the development of CpG-induced MAS. **(A)** Mice received 5 doses of CpG (CpG, ODN 1826, 2 mg/kg), along with daily injections of vehicle (Veh; PBS, 5% DMSO) or VX765 (VX, 100 mg/kg) over 10 days. **(B)** Splenic weight normalised to body weight in mice treated with PBS/PBS, CpG/Veh or CpG/VX (n=5). **(C)** Representative images from **(B). (D)** Plasma levels of ferritin in mice treated with PBS/PBS, CpG/Veh or CpG/VX (n=5). **(E)** H&E staining of spleen in mice treated with PBS/PBS, CpG/Veh or CpG/VX. RP= red pulp, WP= white pulp. White arrows denote changes to normal splenic architecture and perturbations to red pulp. **(F-J)** Plasma concentrations of IFNγ (F), IL-6 (G), IL-10 (H), TNF (I), and IL-18 (J) in mice treated repeatedly with PBS/PBS, CpG/Veh or CpG/VX (n=5). Data represent mean ± SEM. ^*^, P<0.05, ^**^, P<0.01, ^***^, P<0.001 determined by a one-way ANOVA with Tukey’s multiple comparisons test.

### Inhibition of IL-1 receptor signalling with IL-1Ra does not impact the key parameters of MAS

Following on from inflammasome inhibition studies, we sought to test the role of IL-1α/β signalling in this model and its function in the pathogenesis of MAS. Presently, inhibition of IL-1α/β signalling with IL-1Ra (anakinra) along with the use of corticosteroids has proven useful in the clinical treatment of MAS (*34-36, 52, 53*), reviewed in (*54, 55*). As silencing IL-1 signalling has shown some promise clinically, we tested anakinra in animals at a dose of 100mg/kg twice daily (S.C.). Following concomitant treatment of anakinra along with induction of CpG-induced hyperinflammation we examined the main parameters of disease to ascertain the role of IL-1α/β signalling in CpG induced MAS (Figure 4A). Use of IL-1Ra was insufficient to prevent splenomegaly (Figure 4B, C), indicating that IL-1 signalling was not a key driver of spleen enlargement. Secondly, we assessed the ability of anakinra to reduce inflammation in this model by examining plasma ferritin levels. Here, we again observed that IL-1α/β did not drive pathogenesis of MAS, with IL-1Ra treated animals displaying no difference in plasma ferritin levels compared to the placebo treated animals (Figure 4D). Finally, IL-1Ra was unable to prevent CpG-associated splenic architecture disruptions (Figure 4E). Examining plasma cytokine effects, there were no significant differences between CpG-treated mice and those who received IL-1Ra (Figure 4F-J). There was a trend toward a decrease in IL-6, IL-10 and IL-18 (Figure 4G, I and J), indicative of general anti-inflammatory effects expected with IL-1Ra use. Though IL-1Ra treatment is an efficacious treatment for MAS patients, it is not sufficient to prevent CpG-induced hyperinflammation in these animals.

**Figure 4:**
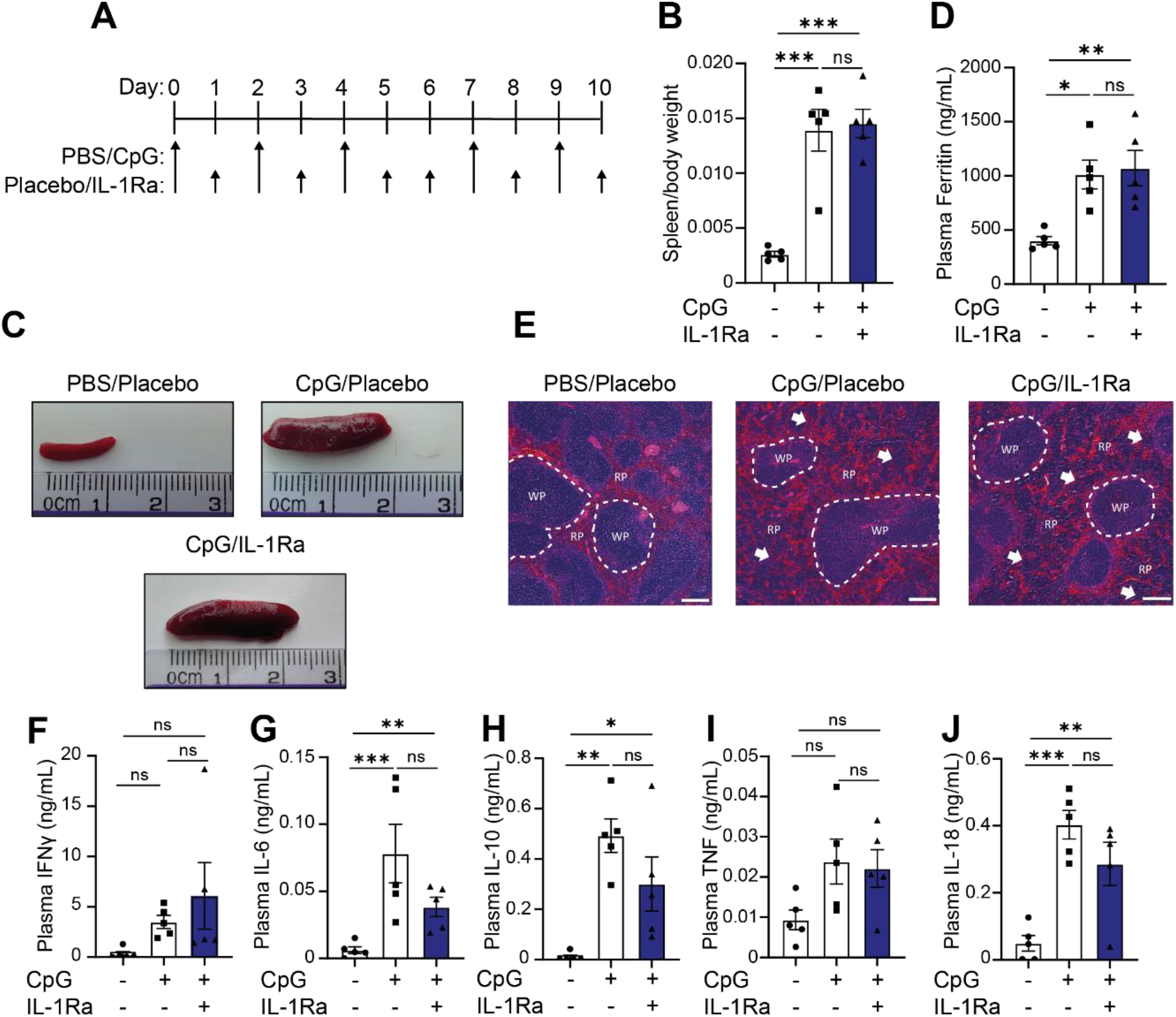
IL-1Ra is not sufficient to prevent onset of hyperinflammation in CpG treated mice. **(A)** Mice received 5 doses of CpG-DNA (CpG, ODN 1826, 2 mg/kg), along with vehicle (placebo) or anakinra (IL1Ra, 100 mg/kg, twice daily). **(B)** Splenic weight normalised to body weight in mice treated with PBS/Veh, CpG/Veh or CpG/IL1Ra (n=5). **(C)** Representative images from **(B). (D)** Plasma levels of ferritin in mice treated with PBS/Veh, CpG/Veh or CpG/IL1Ra (n=5). **(E)** H&E staining of spleen in CpG-treated animals in mice treated with PBS/Veh, CpG/Veh or CpG/IL1Ra. **(F-J)** Plasma concentrations of IFNγ (F), IL-6 (G), IL-10 (H), IL-18 (I), and TNF (J) in mice treated repeatedly with PBS/Veh, CpG/Veh or CpG/Veh (n=5) ^**^, P<0.01, ^***^, P<0.001 determined by a one-way ANOVA with Tukey’s multiple comparisons test.

## Discussion

Inflammasome-derived cytokines are proposed to be involved in hyperinflammation and cytokine storm syndromes (*11, 13, 25-27, 36, 52, 54, 56, 57*) but the role of inflammasomes in the progression of hyperinflammation is not fully understood. Therefore, we sought to identify the roles of inflammasomes in the induction of hyperinflammation using a mouse model of CpG-induced MAS. Here, we show that components of the NLRP3 inflammasome are upregulated following induction of CpG-induced hyperinflammation, and that inflammasomes are critical for the induction of IL-18, a key cytokine involved in the clinical manifestations of MAS (*13, 25, 31*). However, blockade of NLRP3, caspase-1 or IL-1 signalling was not sufficient to prevent the development of several features of hyperinflammatory disease, such as hyperferritinaemia and splenomegaly, suggesting that these traits occur independent of the NLRP3 inflammasome.

This is the first time that inflammasomes have been studied in the development of CpG-induced MAS. However, inflammasomes have already been directly implicated in NLRC4-MAS (*42-44, 58*). Gain-of-function mutations in the NLRC4 inflammasome, leading to aberrant NLRC4 inflammasome activation are characterised as ‘IL-18opathies’, as there is such a strong IL-18 signature associated with these mutations. This pathology is very similar to what is observed in other instances of hyperinflammation, including MAS not caused by mutations in NLRC4. Here, we have demonstrated that in CpG-induced MAS, inhibition of the NLRP3 inflammasome resulted in a reduction in plasma IL-18 levels, indicating that the NLRP3 inflammasome was the source of IL-18 in CpG-induced MAS. Thus, it is possible that the NLRP3 inflammasome could be a source of IL-18 in clinical instances of MAS without NLRC4 mutations. Further evidence for inflammasome activation in patients with MAS is evident as the use of IL-1 signalling blockade has proven efficacious in the management of MAS flares in SJIA and AOSD. High dose IL-1Ra treatment has proven to be effective in patients with refractory MAS (*49, 54, 59, 60*), as well as proving beneficial in the treatment of MAS in sepsis patients (*61*). However, our data did not support the use of IL-1Ra in the context of this mouse model of hyperinflammation, potentially because these animals had no underlying chronic inflammatory disease, infection or existing co-morbidities.

In patients, the IL-18/IFNγ axis is understood to be essential for driving disease pathogenesis, whereas the importance of IL-18 and IFNγ in animal models of MAS is less clear. We show that inhibition of the NLRP3 inflammasome in CpG-induced MAS resulted in a significant reduction in plasma IL-18 levels but did not alter several features of MAS pathogenesis. This supports previous studies which have characterised the effect of IL-18 receptor (IL-18R) inhibition on CpG-DNA induced MAS severity (*32*). Our data demonstrated that IL-18 was not essential for the development of characteristics associated with hyperinflammatory disease including hyperferritinemia, splenomegaly and cytokine storm, mirroring the results observed in response to direct inhibition of IL-18 signalling (*32*). Although IL-18 is thought to be a key cytokine involved in the clinical pathogenesis of MAS, it is not essential in this model. Further, our data demonstrate that in CpG-induced MAS, IL-18 and IFNγ appeared to be uncoupled, where inhibition of the NLRP3 inflammasome was sufficient to reduce plasma IL-18, whilst plasma IFNγ remained elevated. Originally IL-18 was coined as IFNγ inducing factor (IGIF) (*62, 63*), though its role in immunity has become more widely studied (*64, 65*), induction of IFNγ remains a key role for IL-18. Our data indicated that IL-18 is not an essential driver of IFNγ in CpG-induced MAS. Interestingly, IFNγ blockade has been shown to be effective in reducing several of the pathologies associated with hyperinflammation in CpG-induced MAS, including splenomegaly and plasma cytokines (*40*). Furthermore, inhibition of IFNγ in CpG-induced MAS resulted in a reduction in MAS severity in IL-18BP knockout mice (*32*), indicating that IFNγ is an important player in disease pathogenesis. Although not significant, our data suggests that there was a trend towards an increase in plasma levels of IFNγ in animals with reduced IL-18 concentrations, alluding to a change in the signature of the cytokine storm.

Whilst our data suggests inflammasomes and IL-18 play a non-essential role in several manifestations of MAS, it must be acknowledged that our study was performed on young and healthy mice without pre-existing comorbidities. In patients, MAS flares commonly occur with an underlying inflammatory disease, such as SJIA and AOSD (*3, 4, 11, 66*), that have a significant level of basal inflammation which may be crucial to fully recapitulate the disease. Clinically, IL-18 is demonstrated to be a critical cytokine in MAS pathogenesis as: 1) many patients presenting with hyperinflammatory disease present with significantly elevated plasma IL-18 (*56, 67*), 2) plasma IL-18 concentrations correlate strongly with disease severity (*25*), and 3) targeting IL-18 signalling has shown promise as treatment strategy for MAS (*68*). Conversely, our data suggests that IL-18 is not important in CpG-induced MAS in young and healthy animals, since inhibition of NLRP3 or caspase-1 was able to significantly reduce plasma IL-18 levels without preventing the development of MAS. Therefore, we propose that whilst the NLRP3 inflammasome was necessary to produce plasma IL-18 in this model of MAS, there could be intrinsic mechanisms regulating IL-18 dependent inflammation in healthy, young mice meaning that our results have not reflected those observed in patients with hyperinflammatory disease. For example, IL-18BP knockout mice treated with CpG-DNA develop a more severe disease, indicating that endogenous regulation of IL-18 is paramount for controlling disease severity (*25*). Importantly, since we have identified the NLRP3 inflammasome as the source of IL-18 in CpG-induced MAS, targeting the NLRP3 inflammasome in patients with MAS, where IL-18 signalling is crucial, could be an effective treatment to alleviate the disease. Future studies using animals with pre-existing conditions such as obesity, chronic inflammatory disease, infection, or ageing is required to reveal more insight into the involvement of the inflammasome in a more severe disease phenotype that more closely mirrors what is observed in patients with MAS flares.

In summary, we have identified that the NLRP3 inflammasome is activated in CpG-induced hyperinflammation and is critical for enhanced IL-18 production, a key driver of hyperinflammatory disease in patients. However, in the CpG-induced model of MAS, NLRP3 and IL-18 are not required for the characteristic manifestations of hyperinflammation, including splenomegaly and hyperferritinaemia. This study sheds new light on the dynamics of hyperinflammation observed in MAS, suggesting that the NLRP3 inflammasome is essential for IL-18 production but is not a driving factor in the pathogenesis of CpG-induced MAS. Since IL-18 is established as critical in the clinical development of MAS, the NLRP3 inflammasome could be an effective target to treat hyperinflammation and requires further research.

## References

1. D. C. Fajgenbaum, C. H. June, Cytokine Storm. N Engl J Med 383, 2255–2273 (2020).

2. D. Ragab, H. Salah Eldin, M. Taeimah, R. Khattab, R. Salem, The COVID-19 Cytokine Storm; What We Know So Far. Front Immunol 11, 1446 (2020).

3. A. Ravelli et al., 2016 Classification Criteria for Macrophage Activation Syndrome Complicating Systemic Juvenile Idiopathic Arthritis: A European League Against Rheumatism/American College of Rheumatology/Paediatric Rheumatology International Trials Organisation Collaborative Initiative. Arthritis Rheumatol 68, 566–576 (2016).

4. M. Iwamoto, Macrophage activation syndrome associated with adult-onset Still’s disease. Nihon Rinsho Meneki Gakkai Kaishi 30, 428–431 (2007).

5. A. C. Liu et al., Macrophage activation syndrome in systemic lupus erythematosus: a multicenter, case-control study in China. Clin Rheumatol 37, 93–100 (2018).

6. C. Turnquist, B. M. Ryan, I. Horikawa, B. T. Harris, C. C. Harris, Cytokine Storms in Cancer and COVID-19. Cancer Cell 38, 598–601 (2020).

7. R. D. Sandler et al., Diagnosis and Management of Secondary HLH/MAS Following HSCT and CAR-T Cell Therapy in Adults; A Review of the Literature and a Survey of Practice Within EBMT Centres on Behalf of the Autoimmune Diseases Working Party (ADWP) and Transplant Complications Working Party (TCWP). Front Immunol 11, 524 (2020).

8. A. Abdelkefi et al., Hemophagocytic syndrome after hematopoietic stem cell transplantation: a prospective observational study. Int J Hematol 89, 368–373 (2009).

9. A. H. Filipovich, S. Chandrakasan, Pathogenesis of Hemophagocytic Lymphohistiocytosis. Hematol Oncol Clin North Am 29, 895–902 (2015).

10. G. S. Schulert, R. Q. Cron, The genetics of macrophage activation syndrome. Genes Immun 21, 169–181 (2020).

11. C. B. Crayne, S. Albeituni, K. E. Nichols, R. Q. Cron, The Immunology of Macrophage Activation Syndrome. Front Immunol 10, 119 (2019).

12. H. Takada et al., Oversecretion of IL-18 in haemophagocytic lymphohistiocytosis: a novel marker of disease activity. Br J Haematol 106, 182–189 (1999).

13. J. M. Krei, H. J. Moller, J. B. Larsen, The role of interleukin-18 in the diagnosis and monitoring of hemophagocytic lymphohistiocytosis/macrophage activation syndrome - a systematic review. Clin Exp Immunol 203, 174–182 (2021).

14. C. A. Dinarello, Biologic basis for interleukin-1 in disease. Blood 87, 2095–2147 (1996).

15. F. L. van de Veerdonk, M. G. Netea, C. A. Dinarello, L. A. Joosten, Inflammasome activation and IL-1beta and IL-18 processing during infection. Trends Immunol 32, 110–116 (2011).

16. P. Broz, V. M. Dixit, Inflammasomes: mechanism of assembly, regulation and signalling. Nat Rev Immunol 16, 407–420 (2016).

17. F. Martinon, K. Burns, J. Tschopp, The inflammasome: a molecular platform triggering activation of inflammatory caspases and processing of proIL-beta. Mol Cell 10, 417–426 (2002).

18. K. Schroder, J. Tschopp, The inflammasomes. Cell 140, 821–832 (2010).

19. E. Latz, T. S. Xiao, A. Stutz, Activation and regulation of the inflammasomes. Nat Rev Immunol 13, 397–411 (2013).

20. N. Kayagaki et al., NINJ1 mediates plasma membrane rupture during lytic cell death. Nature 591, 131–136 (2021).

21. P. Seckinger, J. W. Lowenthal, K. Williamson, J. M. Dayer, H. R. MacDonald, A urine inhibitor of interleukin 1 activity that blocks ligand binding. J Immunol 139, 1546–1549 (1987).

22. D. Novick et al., Interleukin-18 binding protein: a novel modulator of the Th1 cytokine response. Immunity 10, 127–136 (1999).

23. K. Nakamura, H. Okamura, M. Wada, K. Nagata, T. Tamura, Endotoxin-induced serum factor that stimulates gamma interferon production. Infect Immun 57, 590–595 (1989).

24. H. Okamura et al., Cloning of a new cytokine that induces IFN-gamma production by T cells. Nature 378, 88–91 (1995).

25. E. S. Weiss et al., Interleukin-18 diagnostically distinguishes and pathogenically promotes human and murine macrophage activation syndrome. Blood 131, 1442–1455 (2018).

26. M. Shimizu et al., Distinct cytokine profiles of systemic-onset juvenile idiopathic arthritis-associated macrophage activation syndrome with particular emphasis on the role of interleukin-18 in its pathogenesis. Rheumatology (Oxford) 49, 1645–1653 (2010).

27. K. Mazodier et al., Severe imbalance of IL-18/IL-18BP in patients with secondary hemophagocytic syndrome. Blood 106, 3483–3489 (2005).

28. M. Shimizu, Y. Nakagishi, A. Yachie, Distinct subsets of patients with systemic juvenile idiopathic arthritis based on their cytokine profiles. Cytokine 61, 345–348 (2013).

29. M. Jelusic et al., Interleukin-18 as a mediator of systemic juvenile idiopathic arthritis. Clin Rheumatol 26, 1332–1334 (2007).

30. C. Girard et al., Elevated serum levels of free interleukin-18 in adult-onset Still’s disease. Rheumatology (Oxford) 55, 2237–2247 (2016).

31. T. Shiga et al., Usefulness of Interleukin-18 as a Diagnostic Biomarker to Differentiate Adult-Onset Still’s Disease With/Without Macrophage Activation Syndrome From Other Secondary Hemophagocytic Lymphohistiocytosis in Adults. Front Immunol 12, 750114 (2021).

32. C. Girard-Guyonvarc’h et al., Unopposed IL-18 signaling leads to severe TLR9-induced macrophage activation syndrome in mice. Blood 131, 1430–1441 (2018).

33. C. Gabay et al., Open-label, multicentre, dose-escalating phase II clinical trial on the safety and efficacy of tadekinig alfa (IL-18BP) in adult-onset Still’s disease. Ann Rheum Dis 77, 840–847 (2018).

34. P. M. Miettunen, A. Narendran, A. Jayanthan, E. M. Behrens, R. Q. Cron, Successful treatment of severe paediatric rheumatic disease-associated macrophage activation syndrome with interleukin-1 inhibition following conventional immunosuppressive therapy: case series with 12 patients. Rheumatology (Oxford) 50, 417–419 (2011).

35. N. Bruck et al., Rapid and sustained remission of systemic juvenile idiopathic arthritis-associated macrophage activation syndrome through treatment with anakinra and corticosteroids. J Clin Rheumatol 17, 23–27 (2011).

36. O. Phadke, K. Rouster-Stevens, H. Giannopoulos, S. Chandrakasan, S. Prahalad, Intravenous administration of anakinra in children with macrophage activation syndrome. Pediatr Rheumatol Online J 19, 98 (2021).

37. E. M. Behrens et al., Repeated TLR9 stimulation results in macrophage activation syndrome-like disease in mice. J Clin Invest 121, 2264–2277 (2011).

38. M. B. Jordan, D. Hildeman, J. Kappler, P. Marrack, An animal model of hemophagocytic lymphohistiocytosis (HLH): CD8+ T cells and interferon gamma are essential for the disorder. Blood 104, 735–743 (2004).

39. F. E. Sepulveda et al., Distinct severity of HLH in both human and murine mutants with complete loss of cytotoxic effector PRF1, RAB27A, and STX11. Blood 121, 595–603 (2013).

40. D. K. Gao et al., IFN-gamma is essential for alveolar macrophage-driven pulmonary inflammation in macrophage activation syndrome. JCI Insight 6, (2021).

41. S. W. Canna et al., Interferon-gamma mediates anemia but is dispensable for fulminant toll-like receptor 9-induced macrophage activation syndrome and hemophagocytosis in mice. Arthritis Rheum 65, 1764–1775 (2013).

42. S. W. Canna et al., An activating NLRC4 inflammasome mutation causes autoinflammation with recurrent macrophage activation syndrome. Nat Genet 46, 1140–1146 (2014).

43. C. T. Chear et al., A novel de novo NLRC4 mutation reinforces the likely pathogenicity of specific LRR domain mutation. Clin Immunol 211, 108328 (2020).

44. S. W. Canna et al., Life-threatening NLRC4-associated hyperinflammation successfully treated with IL-18 inhibition. J Allergy Clin Immunol 139, 1698–1701 (2017).

45. J. C. McGrath, E. Lilley, Implementing guidelines on reporting research using animals (ARRIVE etc.): new requirements for publication in BJP. Br J Pharmacol 172, 3189–3193 (2015).

46. R. C. Coll et al., A small-molecule inhibitor of the NLRP3 inflammasome for the treatment of inflammatory diseases. Nat Med 21, 248–255 (2015).

47. W. Wannamaker et al., (S)-1-((S)-2-[1-(4-amino-3-chloro-phenyl)-methanoyl]-amino-3,3-dimethyl-butanoyl)-pyrrolidine-2-carboxylic acid ((2R,3S)-2-ethoxy-5-oxo-tetrahydro-furan-3-yl)-amide (VX-765), an orally available selective interleukin (IL)-converting enzyme/caspase-1 inhibitor, exhibits potent anti-inflammatory activities by inhibiting the release of IL-1beta and IL-18. J Pharmacol Exp Ther 321, 509–516 (2007).

48. M. Sun et al., Treatment with an interleukin-1 receptor antagonist mitigates neuroinflammation and brain damage after polytrauma. Brain Behav Immun 66, 359–371 (2017).

49. P. J. Kahn, R. Q. Cron, Higher-dose Anakinra is effective in a case of medically refractory macrophage activation syndrome. J Rheumatol 40, 743–744 (2013).

50. S. J. Carter, R. S. Tattersall, A. V. Ramanan, Macrophage activation syndrome in adults: recent advances in pathophysiology, diagnosis and treatment. Rheumatology (Oxford) 58, 5–17 (2019).

51. D. B. Kell, E. Pretorius, Serum ferritin is an important inflammatory disease marker, as it is mainly a leakage product from damaged cells. Metallomics 6, 748–773 (2014).

52. S. Ajeganova, A. De Becker, R. Schots, Efficacy of high-dose anakinra in refractory macrophage activation syndrome in adult-onset Still’s disease: when dosage matters in overcoming secondary therapy resistance. Ther Adv Musculoskelet Dis 12, 1759720X20974858 (2020).

53. L. Naymagon, Anakinra for the treatment of adult secondary HLH: a retrospective experience. Int J Hematol 116, 947–955 (2022).

54. P. Mehta, R. Q. Cron, J. Hartwell, J. J. Manson, R. S. Tattersall, Silencing the cytokine storm: the use of intravenous anakinra in haemophagocytic lymphohistiocytosis or macrophage activation syndrome. Lancet Rheumatol 2, e358–e367 (2020).

55. T. A. Gleeson et al., Looking into the IL-1 of the storm: are inflammasomes the link between immunothrombosis and hyperinflammation in cytokine storm syndromes? Discovery Immunology 1, (2022).

56. S. Yasin et al., IL-18 as a biomarker linking systemic juvenile idiopathic arthritis and macrophage activation syndrome. Rheumatology (Oxford) 59, 361–366 (2020).

57. S. M. Vora, J. Lieberman, H. Wu, Inflammasome activation at the crux of severe COVID-19. Nat Rev Immunol 21, 694–703 (2021).

58. J. Barsalou et al., Rapamycin as an Adjunctive Therapy for NLRC4 Associated Macrophage Activation Syndrome. Front Immunol 9, 2162 (2018).

59. P. Mehta et al., COVID-19: consider cytokine storm syndromes and immunosuppression. Lancet 395, 1033–1034 (2020).

60. S. Aytac et al., Macrophage activation syndrome in children with systemic juvenile idiopathic arthritis and systemic lupus erythematosus. Rheumatol Int 36, 1421–1429 (2016).

61. B. Shakoory et al., Interleukin-1 Receptor Blockade Is Associated With Reduced Mortality in Sepsis Patients With Features of Macrophage Activation Syndrome: Reanalysis of a Prior Phase III Trial. Crit Care Med 44, 275–281 (2016).

62. M. Tone, S. A. Thompson, Y. Tone, P. J. Fairchild, H. Waldmann, Regulation of IL-18 (IFN-gamma-inducing factor) gene expression. J Immunol 159, 6156–6163 (1997).

63. Y. Gu et al., Activation of interferon-gamma inducing factor mediated by interleukin-1beta converting enzyme. Science 275, 206–209 (1997).

64. C. A. Dinarello, D. Novick, S. Kim, G. Kaplanski, Interleukin-18 and IL-18 binding protein. Front Immunol 4, 289 (2013).

65. S. A. Ihim et al., Interleukin-18 cytokine in immunity, inflammation, and autoimmunity: Biological role in induction, regulation, and treatment. Front Immunol 13, 919973 (2022).

66. A. Lenert, Q. Yao, Macrophage activation syndrome complicating adult onset Still’s disease: A single center case series and comparison with literature. Semin Arthritis Rheum 45, 711–716 (2016).

67. M. Kawashima et al., Levels of interleukin-18 and its binding inhibitors in the blood circulation of patients with adult-onset Still’s disease. Arthritis Rheum 44, 550–560 (2001).

68. S. Yasin et al., IL-18 as therapeutic target in a patient with resistant systemic juvenile idiopathic arthritis and recurrent macrophage activation syndrome. Rheumatology (Oxford) 59, 442–445 (2020).

