## Supplementary figures for "The NLRP3 inflammasome is essential for IL-18 production in a murine model of macrophage activation syndrome"

**Supplementary Data**

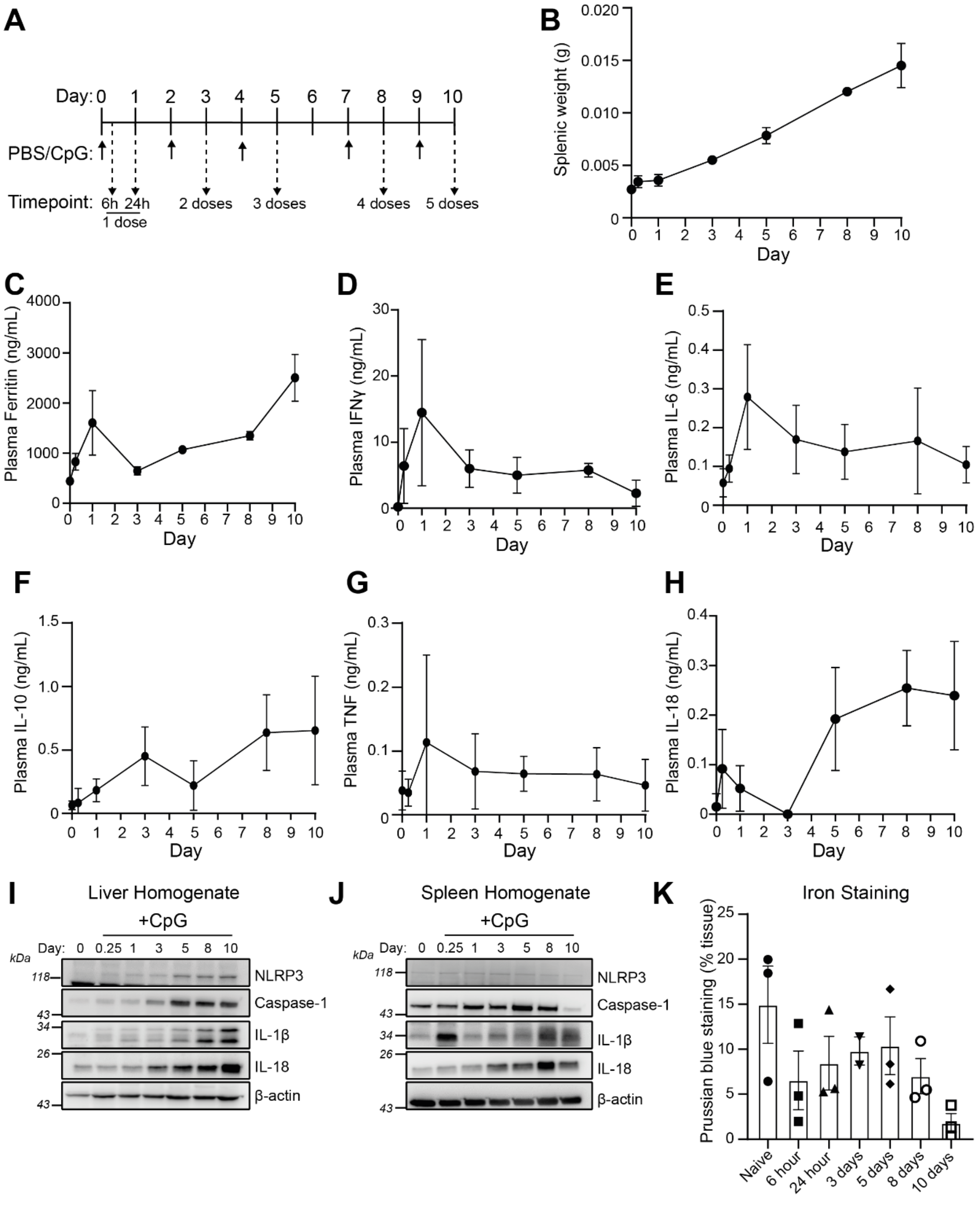

**Supplemental Figure 1: The upregulation of Inflammasome signalling requires multiple doses of CpG-DNA. (A)** Animals received CpG-DNA (ODN 1826, 2 mg/kg) administered via I.P. injection as indicated. **(B)** Splenic weight normalised to body weight over time in mice treated as in (A) (n=3). **(C-H)** Plasma levels of ferritin (C), IFNγ (D), IL-6 (E), IL-10 (F), TNF (G) and IL-18 (H) over time in mice treated as in (A) (n=3). **(F-G)** Western blot analysis of liver (I) and spleen (J) homogenate from mice treated as in (A) of the inflammasome components NLRP3, caspase-1, IL-18 and IL-1β (n=3) **(K)** Iron staining of liver over time in mice treated with CpG-DNA. Prussian blue staining was quantified in Qupath and staining is expressed as a percentage of total tissue area (n=3, apart from day 3, where n=2).
